# A Deep Reinforcement Learning based approach for movement training of neuro-musculoskeletal systems

**DOI:** 10.1101/2021.03.28.437396

**Authors:** Raghu Sesha Iyengar, Kapardi Mallampalli, Mohan Raghavan

**Affiliations:** Indian Institute of Technology, Hyderabad, INDIA

**Keywords:** spinal cord model, movement training, deep reinforcement learning

## Abstract

Mechanisms behind neural control of movement have been an active area of research. Goal-directed movement is a common experimental paradigm used to understand these mechanisms and relevant neural pathways. In this paper, we attempt to build an anatomically and physiologically realistic model of spinal cord along with the relevant circuitry and interface it with a musculoskeletal model of an upper limb, using the NEUROiD platform. The neuronal model (simulated on NEURON) and the musculoskeletal model (simulated on OpenSim) are cosimulated on NEUROiD. We then use Deep Reinforcement Learning to obtain a functionally equivalent model of the supraspinal components and the descending cortical activations feeding into the last-order interneurons and motoneurons. Uniplanar goal directed movement of the elbow joint was used as the goal for the learning algorithm. Key aspects of our work are: (1) Our solution converges naturally to the triphasic response observed in goal directed tasks (2) Gradually increasing the complexity of task helped in faster learning (3) In response to corticospinal inputs, our model could produce movements on which it was not explicitly trained, but were close to the trained movements. Being able to generate movements on which the model was not explicitly trained, implies that the movement repertoire that a biomimetic model needs to learn, could be much smaller than the complete set of movements it can execute. We hope that this will lead to building larger and complex biomimetic systems, one block at a time.

## I. Introduction

The role of cortical areas in processing sensory feedback, movement planning and control is an active area of research. Though simple locomotion has been shown to be possible without any cortical input ([1]) in rats, voluntary reaching and precise positioning tasks, require cortical involvement ([2], [3]). The motor cortex is also implicated with fine motor movement in primates possibly via the lateral motor column ([4], [5]). The mono (mostly excitatory) and oligo (mostly inhibitory, via interneurons) synaptic connection between the cortical cells and motoneurons is well established ([6]–[8]). The spinal locomotor system comprising of all the spinal circuits is controlled via the mesencephalon and diencephalon locomotor region (MLR and DLR) ([9]). The spinal networks are capable of orchestrating stereotypic locomotor pattern by activating specific interneurons and/or motoneurons ([10]).

These spinal networks have been studied in the context of undulatory motion ([11], [12]), flexor-extensor coordination in mammals ([13]), left-right coordination ([14]) and hindlimb-forelimb activation ([15]). Muscle spindle, joints and cutaneous sensory feedback are essential to control movements. Ia sensory afferent fibers respond to the rate of change in muscle length, as well as to change in velocity, while the II sensory afferent fibers respond to muscle stretch ([10]). The roles of Ia, II and Ib fibers in motor feedback and the nature of proprioceptive responses during locomotion has been well studied and modelled ([16], [17]).

However, the exact nature of movement control and movement selection ([18]) by the motor cortex is still unclear, especially in the control of the upper limb in vertebrates, with multiple proposed models. While it is generally agreed that cortical motor activity reflects more than just muscle activity, there is no consensus on what else is encoded and the subset of neurons responsible for the same ([19]). Cortical output from a subset of neurons, residing in caudal M1 that presents cortical output to the alpha motor neurons, commonly called as the corticomotoneurons(CM) has been found to correlate with various movement parameters such as position([20]), velocity([21], [22]), force ([23]), rate of change of torque ([24]) or EMG activity of the target muscle([25]). However, in contrast with alpha motor neurons, the CM neuron activity is correlated to muscle activity only for special classes of movements. Another theory posits that M1 neurons code for temporally extensive movement trajectory fragments ([26], [27]). However it is unclear how or why the cortical circuits come to have neurons with such widely varying tuning properties. It is also unclear how these two types of M1 neurons can interact in order to generate the requisite movement signals in directional movement. Our study aims to understand this question of central importance in the cortical control of movement using learned movements in a simulated movement infrastructure.

The choice of movement models used in this investigation is important. The models required for our purpose should have a spino-musculoskeletal (SMSK) system and a cortical model that can learn to control the SMSK system. Since we know that the cortical networks learn the required control signals for movement control using the incoming proprioceptive and other feedback signals. These inputs are processed in combination with movement intention to generate CM neuron signals that stimulate spinal motor neurons. The spinal motor neurons are additionally influenced by the proprioceptive afferents through the spinal circuits. Thus in order to ensure that the input to and the outputs from cortex are modeled sufficiently realistically, the patterns of proprioceptive signals need to be as realistically as possible. The musculoskeletal anatomy and the spinal reflex networks, in short the spino-musculoskeletal system has a significant bearing on the temporal patterns of proprioceptive signals. This necessitates an anatomically realistic musculoskeletal system coupled to a spinal network.

Since the M1 control circuits and the input proprioceptive afferents are not clearly known, we choose to use an artificial neural network model with the Reinforcement learning(ANN-RL) paradigm to play the role of M1. We further explore the trained ANN-RL to gain insights into the multiple mechanisms by which the different known tuning properties of M1 neurons may arise. In this paper, we build a neuromotor model of upperlimb using NEUROiD ([28]). The model includes the relevant sections of spinal cord, neuronal cell types and spinal networks. A musculoskeletal model of the upperlimb is interfaced to the neuronal model. Both neuronal and musculoskeletal model are cosimulated (refer figure 1). By using NEURON and OpenSim for simulation in the backend, we gain from advances in both these widely supported simulators in the two domains. We then wrap the neuro-musculoskeletal model in a Reinforcement Learning ([29]) framework.

**Fig. 1.**
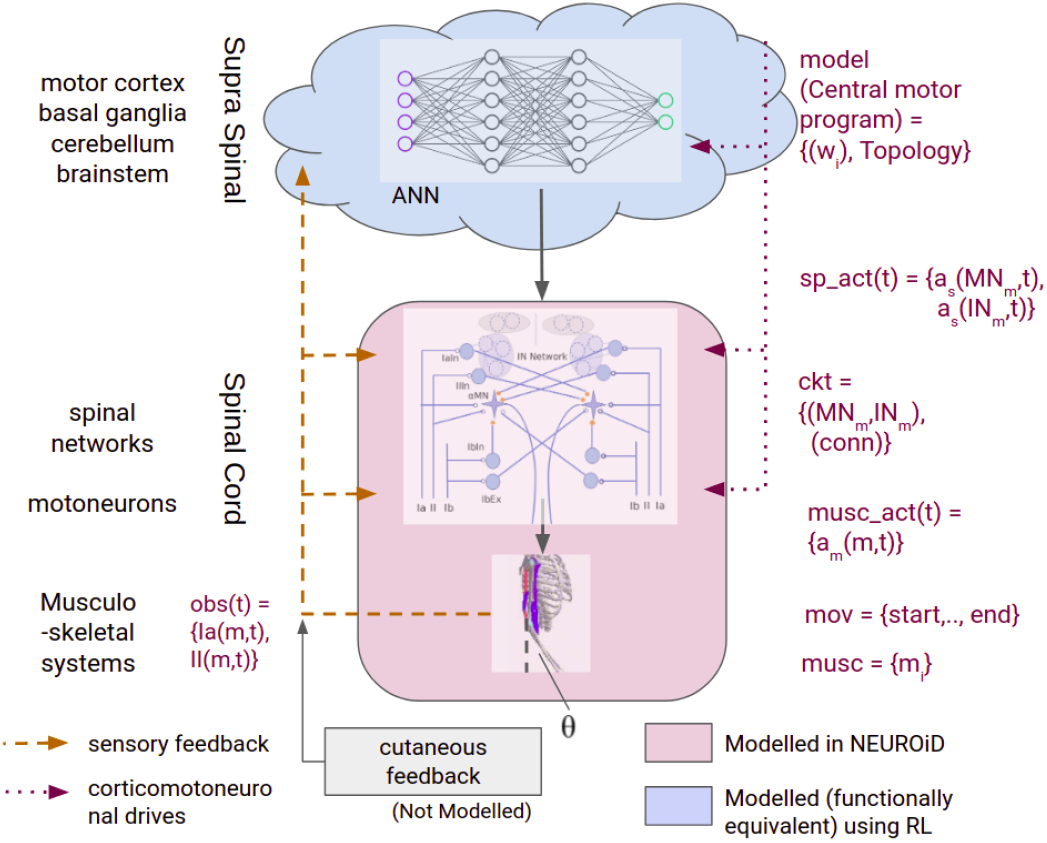
Components of nervous system involved in movement control. The components modelled in NEUROiD and those learnt using Reinforcement Learning are highlighted.

In this work, we show that the policy learnt by the reinforcement learning algorithm is functionally equivalent to the M1 in the context of a specific goal directed movement. The muscle activations from the resultant movement have a close resemblance to physiologically observed triphasic response ([30], [31]). Addition of a small bias to the afferent signals resulted in a change of target position. Thus, mimicing the trajectory selection operation in M1. We further analyzed the ANN obtained as part of reinforcement learning. It was found that the activations of the neurons directly activating the spinal networks correlated well with the movement velocity, mimicing the CM neurons ([21], [22]). Specific other neurons in the ANN were found to encode the trajectory ([26]).

Section II provides more details on the model and the setup. In section III, we describe the results and in section IV, we discuss the significance of our work and its future.

## II. Methods

In this section, we describe the methods for developing a virtual human physiology model that performs a given behavioural task. We first give a formulation for the problem, followed by details of the setup and methodology used.

### A. Problem Formulation

Figure 1 shows the problem formulation for the current work. we are interested in producing a movement on the virtual limb, where the movement (*mov*={*start*,‥,*end*}) is defined by a start position of the limb, a desired target position and possibly zero or more intermediate positions. A set of muscles (*musc*={*m_i_*}) which are known to be responsible for causing the desired movement are activated by a time series of activations (*musc*_*act*(*t*)={*a_m_*(*m,t*)}) provided by the last-order interneurons and motoneurons of the relevant spinal networks. These spinal networks (*ckt* = {(*MN_m_*, *IN_m_*),(*conn*)}) are comprised of the set of motoneurons, interneurons and a connectivity matrix. The supraspinal centers involved in generating the desired movement (*mov*) are grouped into a ‘‘central motor program” (*model(central motor program)* = {*W_i,topology_*}). A Central Motor Program (CMP) provides the necessary activations (*sp*_*act*(*t*)={*a_s_*(*MN_m,t_*), *a_s_*(*IN_m_*,*t*)}) to the spinal (*ckt*) to cause a specific movement. Our aim now is to obtain a functionally equivalent representation for the CMP, using machine learning techniques.

### B. NEUROiD

Computational models aimed at movement understanding are focussed primarily on either the neuronal ([32]–[34]) or musculoskeletal components ([35], [36]). Some models include both these components. However, in these models ([37]–[40]), the individual components are built and do not take advantage of the advances in respective fields. In some models, the neuronal components are detailed, while the musculoskeletal models are relatively simplistic ([41], [42]). A model with detailed spinal circuits was built to demonstrate wrist flexion ([40]). However, the pools of interneurons and motorneurons were modelled as simplistic sigmoid function. Some models have also attempted to include the effect of cortical inputs ([37], [39]). [39] showed among other things, that interpolation of learnt solution was sufficient to perform previously unseen tasks. AnimatLab([43]) allows its users to build biologically realistic neural systems and biomechanical muscle models.

NEUROiD ([28]) is a spinal cord model integration and simulation platform. It allows neuronal models and musculoskeletal models to be cosimulated. Neuronal models are simulated using NEURON ([44]) and the musculoskeletal models are simulated using OpenSim ([45]). NEUROiD allows a choice of model components from both the domains in a modular fashion, deriving advantage from advances in both the domains. NEUROiD models are inherently multiscale (cellular, network and system) and multidisciplinary (anatomy, physiology and biomechanics) in nature. We use NEUROiD to model and simulate the spinal and musculoskeletal components. Please refer to [28] and [46] for more details on NEUROiD.

### C. Simulation Setup

In this section, we describe the individual components (refer figure 1) in our model.

1. *Musculoskeletal Model:* We assume a simple uni-planar movement of the elbow joint. Thus, the movement we are interested in is {*start* = *0.17 rad*, *end* = *0.43±0.05 rad*} (refer to *θ* in figure 1). The relevant set of muscles for this movement are identified to be {*biceps, brachialis, triceps*}. We used the OpenSim musculoskeletal model of the upper limb based on the Arm26 model ([47]). It consisted of 6 muscles, namely Biceps (2 muscle heads), Triceps (3 muscle heads), and Brachialis. The shoulder joint was constrained while the elbow joint could perform uniplanar movements. The musculoskeleton was in upright position in our experiments. Detailed characterization of this model on NEUROiD can be found in [48]. A snapshot of the musculoskeletal model visualized in OpenSim can be seen as inset in figure 2.
2. *Spinal Network:* The muscles are grouped into flexors and extensors. A timeseries of activations for these muscle groups {*a_m_*(*flex, t*), *a_m_*(*ext, t*)} are obtained from the spinal network. The spinal network is built on the NEUROiD platform by adding the model definitions at the micro, meso and macro levels of the anatomical, physiological and biomechanical components of the spinal cord model. Here, scales micro, meso and macro loosely translate to cellular, network, and system, respectively

a. *System:* We built models for the C5 and C6 sections of the spinal cord. The various cell groups and laminae boundaries were obtained from [49]. We then extended the 2D laminae contours to create 3D regions where the cells were placed. All spinal cord sections were aligned using the central canal region. The lengths of the spinal cord sections were obtained from [50]. A snapshot of the C5 and c6 regions as visualized in NEUROiD can be seen as inset in figure 2.
b. *Network:* A functionally homogenous group of cells present in a specific region forms a cellgroup in NEUROiD. Motoneuron and interneuron cellgroups were placed in appropriate laminae regions based on their cytoarchitectonic localization available from past studies ([49]). We modelled neuronal cell groups involved in controlling the biceps (bi), brachialis (bra) and triceps (tri) muscles. Various pathways in the spinal cord from the literature (summarized by [10]) are defined based on the connectivity between the afferents from a given muscle, its synergist and antagonist motoneurons. NEUROiD uses a unique muscle synergy based circuit generation technique. Using this, we modelled the monosynaptic Ia excitation, reciprocal Ia inhibition, polysynaptic II excitation and Ib inhibitory circuits. Cutaneous inputs were not considered as the tasks that were being modelled did not seem to be influenced by them. Thus, the spinal network used in our model is of the form:

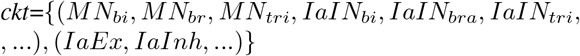
c. *Afferent and Efferent interfaces:* The efferent activation to each muscle is derived from the axonal membrane potential of the corresponding motoneuron ([51]). The spike train derived from the axonal potential is modelled as the sum of a series of time-delayed delta functions.

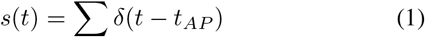

*s*(*t*) is the superposition of all the action potentials occurring at time *t_AP_*. A normalized efferent activation (in range [0,1]) is sent to OpenSim musculoskeletal model. The OpenSim model returns the joint angle, length of muscles, rate of change of length of muscles and the force generated by the muscles after every simulation step. The firing rates of proprioceptive feedback were calculated based on [17]:

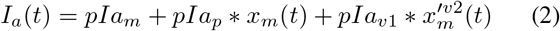

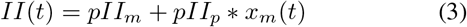

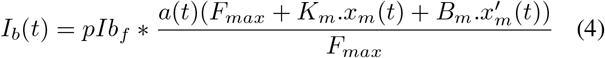 Here, *I_a_*(*t*), *II*(*t*) and *Ib*(*t*) are the instantaneous values of the *a*, *II* and *Ib* afferent firing rates (in PPS). *pIa_m_* (=8OPPS) is the firing rate constant for Ia. *x_m_*(*t*) is muscle stretch in mm, 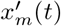 is muscle stretch velocity in mm/s. *pIa_p_* (=2) is length change constant, *p*_*υ*1_ (=4.3) and *υ*2 (=0.6) are velocity constants. *pII_m_* (=5PPS) is the firing rate constant and *pII_p_* (=8) is length change constant for II. *pIb_f_* (=200) is the force constant, *K_m_* (=56.3 kN/m) is muscle stiffness, *B_m_* (=2.81 kNs/m) is damping. *F_max_* is the maximum isometric force of the muscle. *a*(*t*) is the muscle activation (normalized to be in range [0,1]).
d. *Cellular:* Motoneuron cell template in the hoc format was automatically generated in NEUROiD based on the parameters specified. Since these parameters for motoneurons and interneurons are mostly unknown in humans, we use hypothesized parameters based on studies in other mammals. The parameters for the motoneuron cell model are derived from [34]. The soma diameter was set to 80±2.5μm, matching with the values for a S-type motoneuron ([51]). The motoneuron soma membranes included sodium channels, a delayed rectifier potassium, calcium-activated potassium, N-type and L-type calcium currents. The initial segment membrane included fast and persistent sodium channels as well as delayed rectifier potassium currents ([52]). Axons and dendrites were modelled based on results from [32]. We modelled 169 alphamotoneurons, 60 Ia afferents, 60 II afferents and 400 interneurons in each segment of the spinal cord ([34]).
3. *CMP Model:* The spinal network receives its input from the locomotor regions (MLR and DLR) in the brain stem and also the afferent sensory feedback. There is a lot of progress in understandanding the functional roles of various supraspinal regions and their interplay in the context of movement ([53]). Many models of these components have also been built based on the studies. However, a comprehensive framework encompassing all these components, known pathways and their influence on spinal networks in the context of movement is not yet mature. Since spinal cord modelling is the primary focus of NEUROiD, we approach this problem by obtaining a *functionally equivalent* model of the supraspinal components. In this work, we use reinforcemnt learning to obtain the model. The model is approximated by an artificial neural network (ANN) that receives the afferent signals as input and generates the signals (which are equivalent of descending drive signals) to facilitate movement. The CMP is modelled as a trained set of weights (learnt during training) and a topology of the ANN. Thus, *CMP* = {*W_i_*, *Topology*}. The activations to the spinal cord are obtained as a timeseries of flexor and extensor network activations. Thus, the spinal activations from the CMP are modelled as:

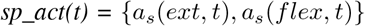 It is known that for a given cortical stimulation, the observed muscle activations vary depending on the state of the joint ([54]). Such state dependent response implies our model of CMP should have access to the afferents and generate (a time series of) patterned input to the spinal networks. It is also apparent that the (functionally equivalent) CMP model should interact with the musculoskeletal model continuously in a closed loop. Hence, We chose reinforcement learning to obtain a model of the CMP.

**Fig. 2.**
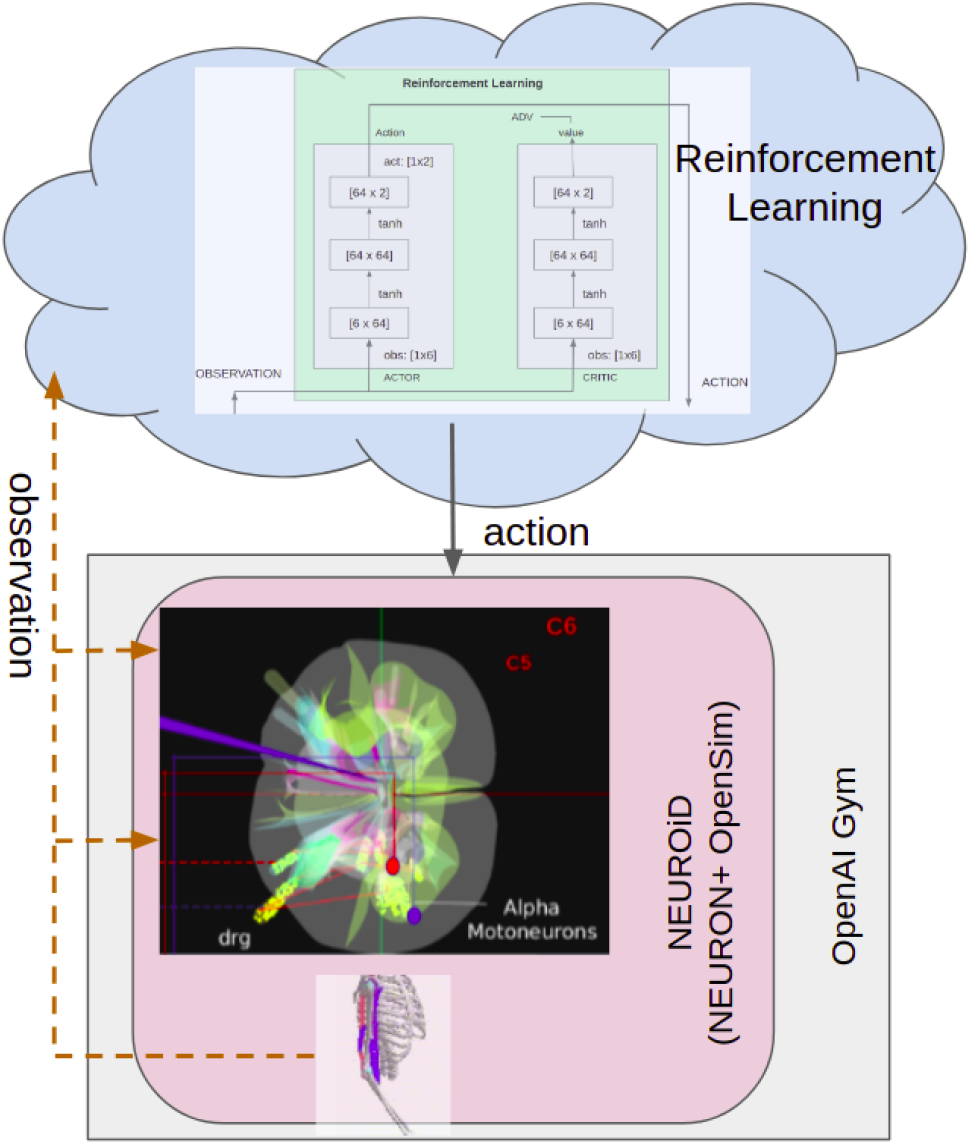
Spinal and musculoskeletal models were built on NEUROiD. Spinal networks were simulated on NEURON and musculoskeletal models were simulated on OpenSim. The NEUROiD (NEURON+OpenSim) model was encapsulated as an OpenAI Gym environment. Reinforcement Learning was used to learn an optimal policy to approximate the Central Motor Program for a specific movement task.

### D. Reinforcement Learning Setup

Reinforcement Learning ([29]) contemplates the problem of an agent interacting with an environment and taking actions with the goal of accumulating maximum expected reward from the environment. Deep Reinforcement Learning uses deep learning for representation learning, but uses the cumulative reward to guide the learning. The neuromusculoskeletal model described in the previous section, is co-simulated on NEURON and OpenSim using NEUROiD. However, in this work, our goal is to identify a functionally equivalent model of the CMP that can be used in performing a goal directed movement of the elbow joint. For this, we encapsulate the NEUROiD (NEURON+OpenSim) model in a reinforcement learning environment (refer figure 2). OpenSim models have been used to develop a controller to achieve walking or running on a physiologically plausible 3D human model ^1^. The contributors for this challenge extensively used Deep RL^2^ to solve this problem. However, all these solutions were restricted to identification of the muscle activations needed to achieve a particular movement and do not model the spinal or supraspinal influences.

We used the OpenAI Gym ^3^ toolkit to create a custom environment with a unified interface to encapsulate the NEUROiD model. OpenAI Gym offers many inbuilt environments to perform research on various reinforcement learning algorithms. It also allows creating custom environments ^4^ so that researchers can use the framework to apply reinforcement learning on problems in various domains. We extended the OpenAI Gym class and encapsulated the NEUROiD environment in that. The observation space of this environment was the Ia, II and Ib afferent firing rates from the muscles. The action space was set to be a vector of two normalized (in the range [0,1]) values that were used to activate the flexor and extensor spinal networks (i.e., *sp_act(t)*).

A brief summary of some of the key methods we implemented as part of the wrapper is given below:

1. *Init:* We initialize the NEUROiD (NEURON+OpenSim) environment in this method along with the observation and action space of the model.
2. *step:* Here, we perform a single step interaction with the environment by taking an action, calculating the reward and reading the observations from the environment:

a. *_take_action:* The activation from the CMP (modelled by the RL policy) is used to update the current stimulation amplitude of the iclamp object stimulating the flexor and extensor network. The muscles in our model are activated based on the corresponding alphamotoneuron firing.
b. *_calc_reward:*: Various components of the reward are shown in figure 3. The dist reward component is high when the model is closer to the desired target position. In our experiments, the goal was set to move the elbow joint to an angle of 0.43 rad from an initial angle of 0.17 rad. Further, we specify that the angle should be held at 0.43 ±0.05 rad for a specified duration. The stay reward component is high when the elbow remains in the target position for the specified duration. The speed reward component ensures that the desired final state is reached as soon as possible. The negative angle penalty is a negative reward that ensures that the elbow angle does not become less than zero. This is a physiologically unrealistic state and adding this component to the reward ensures that the policy learns not to move the elbow joint towards negative angles. Some of these reward components may be thought to be analogous to the error correction and learning functions in cerebellum and basal ganglia ([55]).
c. *_next_observation:* The normalized afferent Ia, II and Ib firing rates of the flexor and extensor form the observation space. These are obtained from the NEUROiD environment.
3. *reset:* This function is to reset the state of the NEUROiD environment. This will be called every time an episode is completed. We define an episode to be complete if either the goal is achieved or the simulation has run for a set amount of time.

**Fig. 3.**
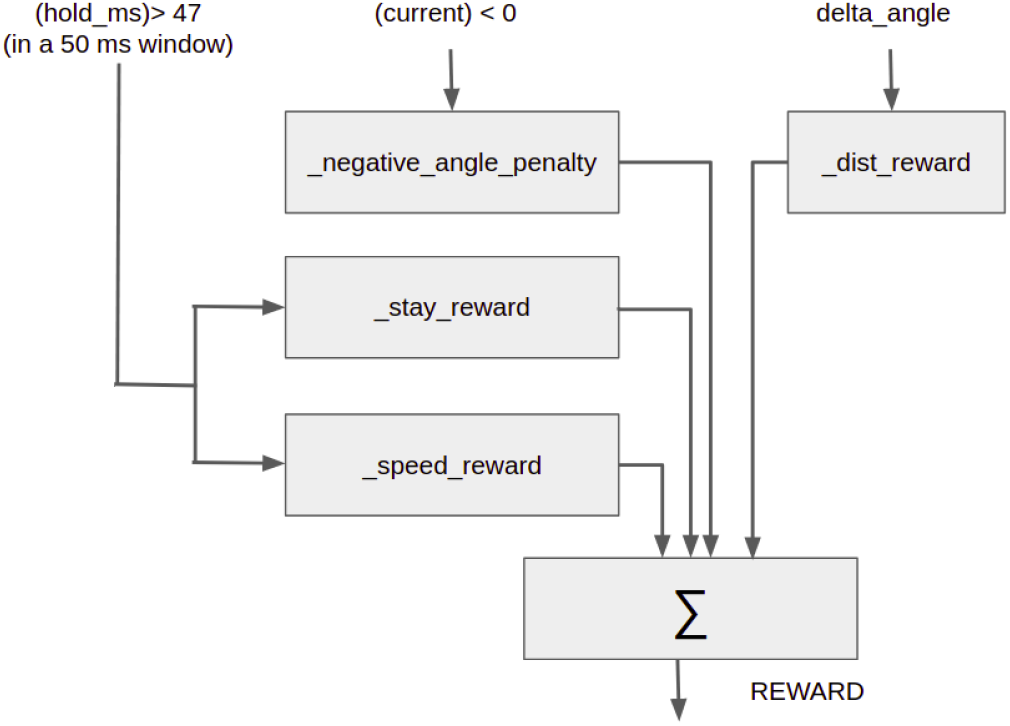
Components of the reward function used in Reinforcement Learning

### E. Training

We used fixed timestep integration in NEURON. The timestep for simulation (dt) was set to 0.025 ms. This is the default value for timestep in NEURON and is necessary to simulate the sub-millisecond dynamics of neuron cell. Open-Sim simulator also runs at 0.025ms timestep. The activations to the muscle model was calculated every 1 ms and sent to OpenSim model using socket communication. This is justified as the muscle response times are atleast of the order of 10’s of ms ([56]). The Ia, II and Ib afferent firing rates are also calculated every 1 ms and sent back to the NEURON model. The Reinforcement Learning action space, observation space and policy are updated every 20 ms.

We used the Proximal Policy Optimization ^5^ to obtain the optimal control policy. PPO uses a novel clipped surrogate objective to search for the optimal policy. This ensures the stability of TRPO ([57]), while reducing the complexity. We use an A2C (Advantage Actor-Critic) ([58], [59]) which uses two separate Actor and Critic networks. The Critic approximates the value function while the Actor estimates the distribution of the action space. The expected value is estimated using a Generalized Advantage Estimation ([60]), which is then used to calculate the advantage of value estimator in current state. We use PPO in an Asynchronous A2C setting (also called A3C). For the purpose of demonstration in this paper, we chose the actor and critic, each to be a two layered fully connected network with 64 neurons in each layer (figure 4). A tanh activation layer is used to model the non-linear dynamics of the system. We used the stable-baselines3^6^ implementation of RL algorithms using pytorch.

**Fig. 4.**
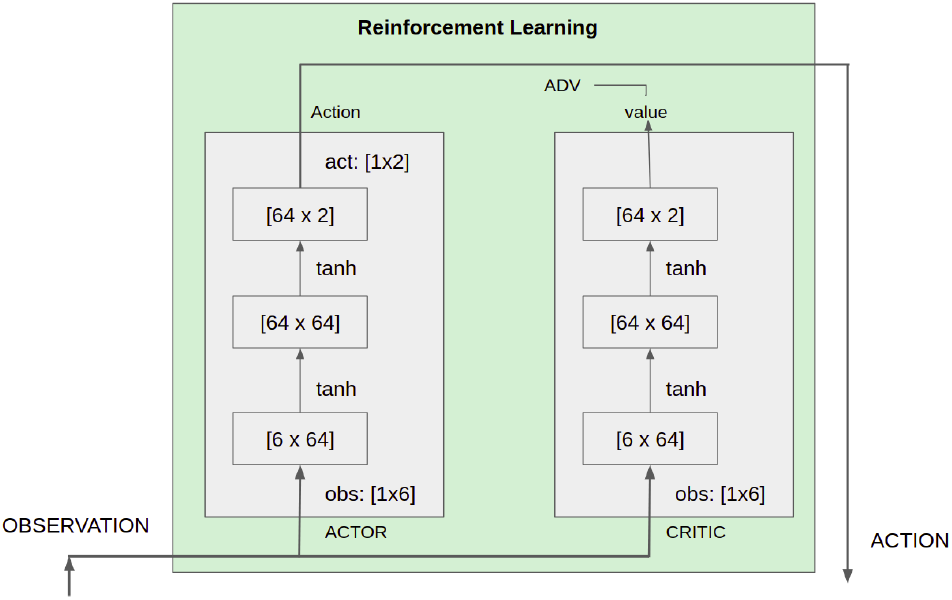
The ANN consisted of separate actor and critic networks. The observation (afferent firing) space was a (1×6) dimension vector. Two hidden layers of dimension 64 were used with tanh layers to model non-linear dynamics. The action (activations to spinal network) space was a ()1×2) dimension vector.

We used a reward discount factor (*γ*) of 0.99 and λ for generalized advantage estimation was set to 0.95. The clip range (*∊*) for clipping the surrogate function was set to 0.2. Adam optimizer with learning rate of 0.0003 was used for learning.

The learning of the functionally equivalent CMP network was attempted with two different configurations - openloop and closeloop. The latter is an upper limb system with the proprioceptive reflexes and the attendant neural circuits intact, while in the former, the proprioceptive afferents back into the spinalcord are disconnected. The openloop training was performed with tasks of gradually increasing complexity and we used the trained openloop weights to initialize closedloop training. This technique, where a machine learning algorithm is trained with tasks whose complexity is gradually increasing is known as Curriculum Learning ([61]). Such techniques have also been applied in robotics and are known to improve speed of convergence and help in generalization. It may be noted that during open loop configuration, the proprioceptive afferents are still available for observation and learning, but the spinal circuits do not have access to them as the reflex loops are open.

## III. Results

In section III-A, we show that the reinforcement learning algorithm was able to learn the necessary policy to achieve a specific goal directed movement, both in openloop and closed loop setup. In section III-B, we show that the learnt policy has a close resemblance to triphasic response ([30], [31]), which is a well known physiological response observed in goal-reach tasks. In section III-C, we show that constant descending drive to the CMP is equivalent to trajectory selection. In section III-D, we analyze the CMP ANN and study its equivalance to M1 in the context of a goal reach task.

### A. Goal reach task with fixed target

1. *Openloop:* Figure 5 shows the plot of average reward Vs number of steps of training in both openloop (deafferentiated) and closeloop (with afferent input) configurations. We start the training with less stringent goals and increase the complexity gradually. In the first experiment (in figure 5 (a)), the initial angle of elbow was set to 0.174 rad and the goal of learning was to hold the elbow at 0.43 ±0.05 rad for at least 15 ms in a window of 30 ms (ext-II-15/30). We then gradually introduced other afferent signals into the observation space and also made the goal more complex. In the next experiment, we add the II afferents flexor also into the observation space and set the goal of learning to hold the elbow at 0.43 ±0.05 rad for at least 30 ms in a window of 40 ms (extflex-II-30/40). We observe that the average reward in this case was slightly lesser than the previous case. However, when we introduce the Ia afferents into the observation space keeping the same goal (extflex-IIIa-30/40), we find that the average reward increases and gets better than the first experiment (ext-II-15/30). II and Ia afferents carry the information about muscle stretch and rate of change of muscle length respectively ([10]). Hence, it makes intuitive sense that giving diverse information about the system helps in learning. When we further add the Ib afferent firing into the observation space and make the goal even tougher (extflex-IIIAIb-47/50), we find that even in this case, the average reward continues to increase with number of steps of training. A typical plot of the actual rewards obtained during the training for an openloop experiment is shown in figure 6. The x-axis here represents the number of steps of learning and the y-axis represents the accumulated reward in an episode. The spikes in the plot correspond to the episodes when the reward was high, indicating that the various components of reward (refer figure 3) were achieved during training.
2. *Closedloop:* Figure 5 (b) shows the plot of average reward Vs number of steps of training in the closed loop setting. We observe that the average reward in the closed loop setting also increases as the training progresses. This improvement in reward was not seen if we performed the close loop training without initialization from the weights trained using open loop. Further, in the closed loop case, we first trained with a relatively easy target as shown in figure 5 (extflex-IaIIIb30/50-CL) and then used the trained weights from this, to train on a tougher target (extflex-IaIIIb-40/50-CL).

**Fig. 5.**
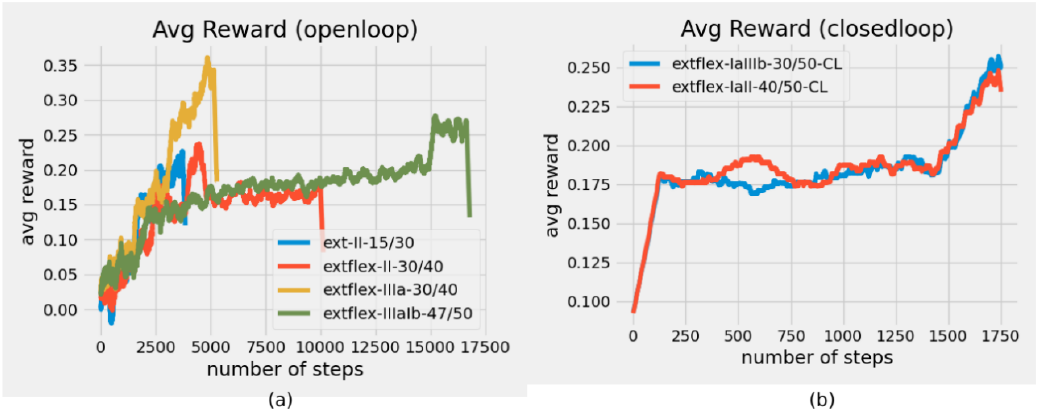
Plot of reward Vs the number of steps of reinforcement learning. The reward is averaged over 100 steps. (a) This is an open loop setup with the NEURON solver running at 0.025 ms, OpenSim muscles are activated every 1 ms and action space in the Open AI gym environment is updated every 20 ms. The plot shows the average reward for slightly different observation spaces and goals. In the above figure, the plots are labelled as follows: ext-II15/30 indicates the experiment where we used the afferent II fibers from the extensor as our observation space. We set the goal of learning to move the elbow joint to an angle of 0.43 rad from an initial angle of 0.17 rad. Further, we specify that the angle should be held at 0.43 ±0.05 for at least 15 ms in a window of 30 ms. extflex-II-30/40 indicates the experiment used the afferent II fibers from the extensor and flexor as our observation space. The target was same for all experiments, however we specify that the angle should be held at 0.43 ±0.05 for at least 30 ms in a window of 40 ms. The observations are used by the Reinforcement Learning algorithm to calculate the state of next action space. The action space consisted of the current amplitude of the two iclamp objects used to stimulate the spinal networks. (b) This is a closed loop setup with the NEURON solver running at 0.025 ms. The plots are labelled as follows: extflex-IIIa-30/50-CL indicates the experiment where we used the afferent II and Ia fibers from the extensor as our observation space. We set the goal of learning to move the elbow joint to an angle of 0.43 rad from an initial angle of 0.17 rad. Further, we specify that the angle should be held at 0.43±0.05 for at least 30 ms in a window of 50 ms, in a closed loop setting.

**Fig. 6.**
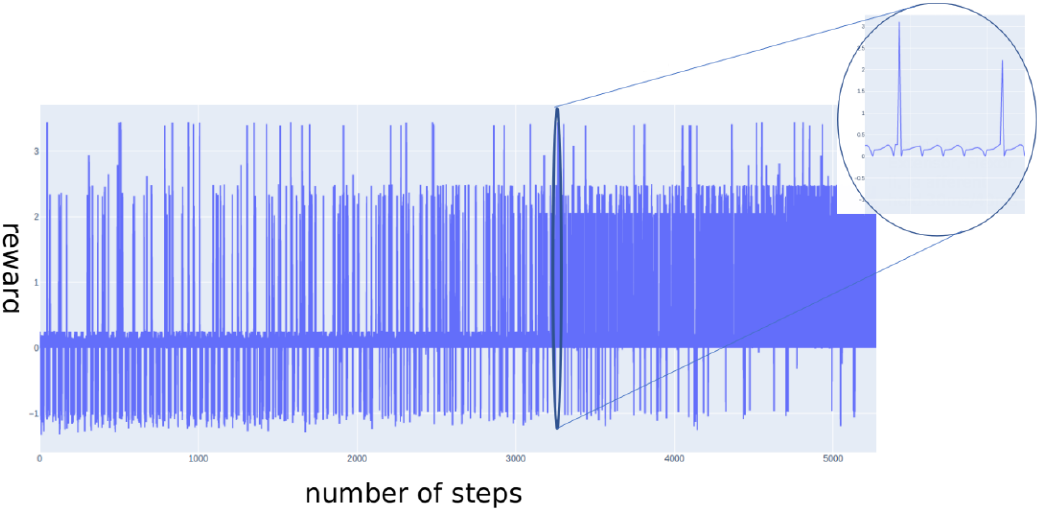
Plot of typical reward Vs the number of steps of reinforcement learning. A small portion of the plot is zoomed in to show the reward for multiple episodes. We observe that (1) the number of trials for which the reward becomes negative decreases as the training progresses. (2) As the training progresses, the algorithm explores the action space and converges towards actions that result in high reward for a larger fraction of episodes.

### B. Muscle activation for goal reach task mimics triphasic response

1. *Reach target from a fixed starting position:* Figure 7 (a) (solid line (S)) shows the force generated by the extensor and flexor muscle groups as observed during the closed loop training after about 12800 steps. The magnitude of the generated force is obtained from OpenSim during simulation. This force is equivalent to convolution of the twitch response of individual muscle and the spike train derived from the axonal membrane potential of the corresponding motoneuron. This can be considered as a proxy for the EMG signal generated by the muscles during contraction ([62]). Figure 1 from [30] shows the typical triphasic pattern observed in EMG during a goal directed task. We observe that the Reinforcement Learning algorithm has learnt to generate a muscle activation pattern similar to the triphasic response. Further, when the velocity of such goal directed movement is increased, the amplitude of the bursts is known to increase with a decrease in duration of each burst ([63], [64]). Figure 7 (a) shows the plot of muscle force after 18900 (dotted line (D)) steps of training. We observe that as training progresses, the amplitude of initial flexor activation becomes larger and the peak extensor activation occurs earlier for “braking”. The second flexor activation for “clamping” is reduced in amplitude as seen in measured physiological response. Since we reward the algorithm if it reaches the goal quickly (refer *_speed_reward* in figure 3), the algorithm naturally learns this physiologically observed behaviour.
2. *Reach target from an arbitrary starting position:* we further train the model by letting the initial angle be a uniform random number between 0.17 rad and 0.7 rad. Figure 7 (b) shows the plot of force generated in the flexor and extensor muscle group with two different initial angles (0.17 and 0.7 rad). We observe that the model has learnt the activations necessary to move the elbow to the specified target angle (0.43 rad) from any initial angle within a range. Such state dependent activation of muscles via cortical stimulations have been observed ([54]). In figure 8 (a), we see that the elbow angle moves towards a desired target angle when the initial angle was set to four different values in the range 0.17 rad - 0.7 rad. A snapshot of the neuromuscular model during the goal-reach task can be seen in figure 8 (b).

**Fig. 7.**
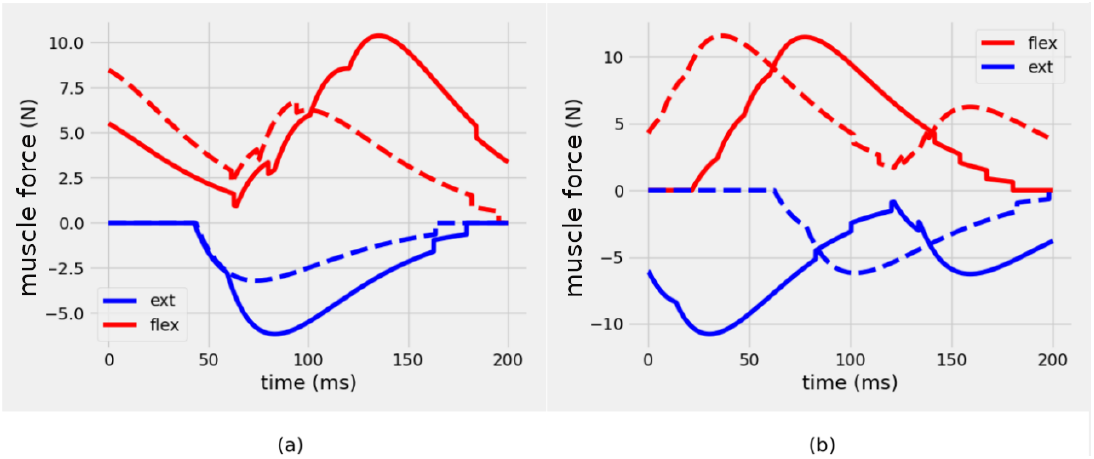
(a) Force generated by the flexor and extensor muscle groups after 12800 (solid line (S)) and 18900 (dash line (D)) steps of training. This force is the result of a time delayed summation of the muscle twitch (i.e., convolution of the twitch response with the spike train obtained from the motoneuron firings). We observe that the initial magnitude of flexor force in (D) is larger than what was observed in (S). The peak of extensor force in (D) is also seen to occur slightly earlier compared to what was observed in (S). The peak of the second activation of flexor in (D) occurs much earlier than in (S). All these observations indicate that the algorithm is learning to reach the goal earlier and hold for longer as the training progresses. (b) Shows the triphasic response observed in the flexor and extensor muscles during training after 18600 steps, when the initial angle is set to 0.7 rad (solid line (S)) and 0.17 rad (dotted line (D)). The target angle is set to 0.43 rad in both cases. We observe that the policy has learnt to activate the flexor and extensor muscles appropriately to reach the goal from both directions.

**Fig. 8.**
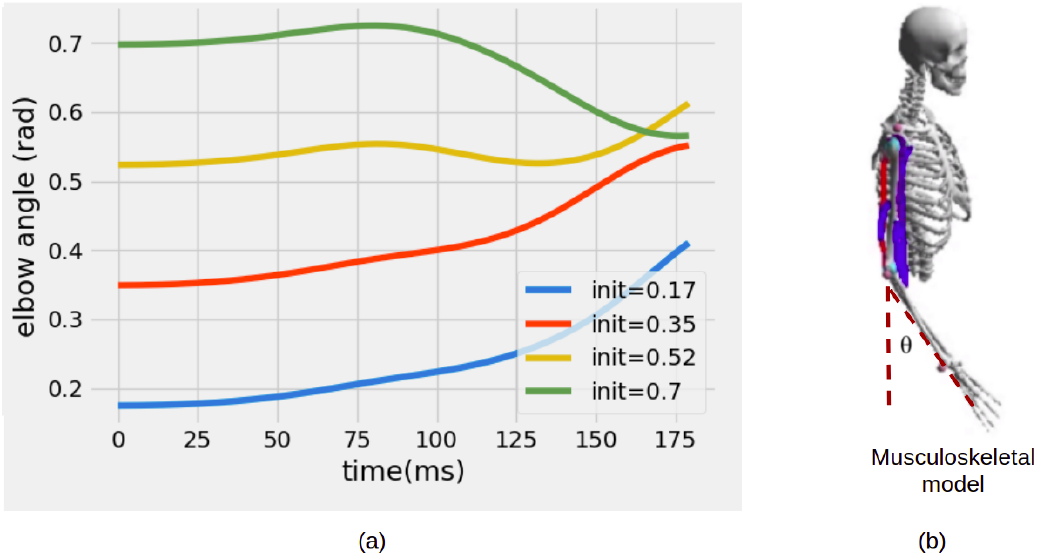
(a) Plot of elbow angle for four different initial angles in range 0.17 rad - 0.7 rad (b) A snapshot of the musculoskeletal model during the goal reach task

**Fig. 9.**
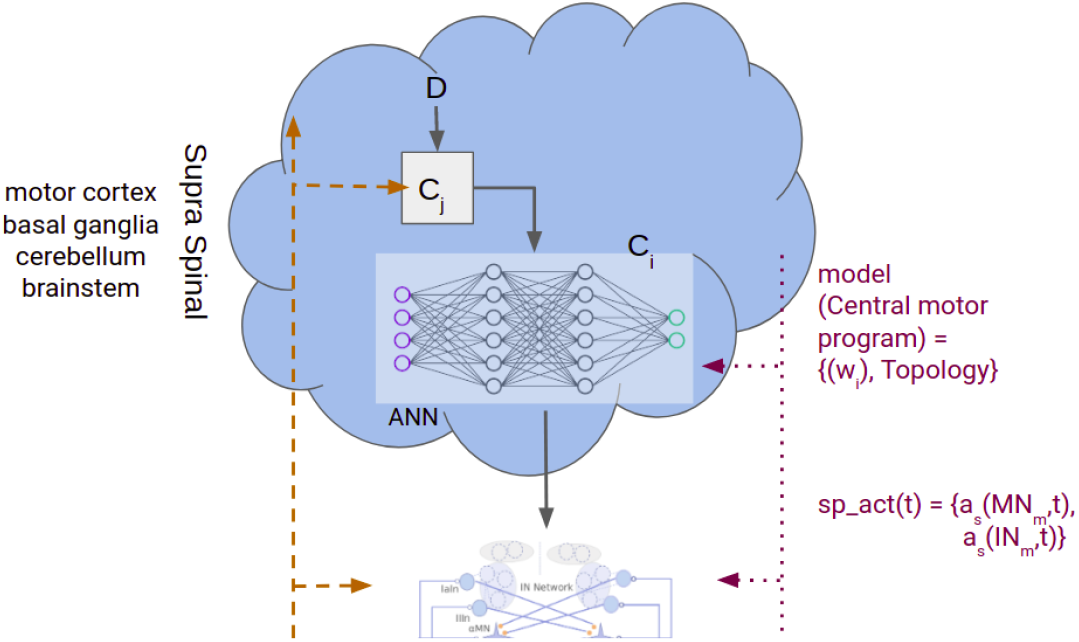
The CMP could be thought to be a set of subcomponents (*C*_1_,*C*_2_,*C*_3_,*C*_i_,*C*_j_,‥*C*_n_). Under special conditions, it may reduce to the already learnt CMP

### C. CMP as compositionality of subcomponents

The CMP identified by Reinforcement Learning is able to provide the necessary time dependent stimulations to the spinal network to achieve the desired movement. We may assume that the learnt CMP (*C_i_*) is a subcomponent within the CMP. We may consider another subcomponent (*C_j_*), that receives the afferent input and higher order descending drive input (*D*). If we assume that *C_j_* does not alter the afferent input before passing it on to *Ci*, then the CMP reduces to the already learnt *C_i_*. It was proposed that neural learning systems use a reasonable approximation of already learnt tasks when a previously unknown task is given ([65]). In [39], a model of spinal movement learning circuit was built. It was shown that interpolation between learnt models was sufficient to execute tasks which it was previously not exposed to. In our work, we propose that the subcomponent *C_j_* plays the role of modulating the afferent signals using the descending drive input (D). In the absence of descending drive input, the CMP reduces to the trained *C_i_* that was obtained using RL. By changing the descending drive, we show that a different target location is achieved. This could be thought of to be equivalent to converging to the same target in a transformed coordinate axes. Such descending connections from brainstem or cortical areas are known to play a role in movement ([11], [66]) or in initiation of Central Pattern Generators ([5]).

Figure 10 (a) shows the plots of the elbow angle when the descending input (marked as (Δ) in figure) is a constant bias. The figure shows these plots for two different initial angles, 0.17 rad (solid line (S)) and 0.7 rad (dashed line (D)). We observe that this effects a change in the target angle in a single-joint, uni-planar setup without any further training on the already learnt CMP. Thus, a constant bias descending drive results in a transformation in the coordinate space, resulting in moving the target to a different location. In figure 10 (b), we observe the same behaviour in a closed loop setting.

**Fig. 10.**
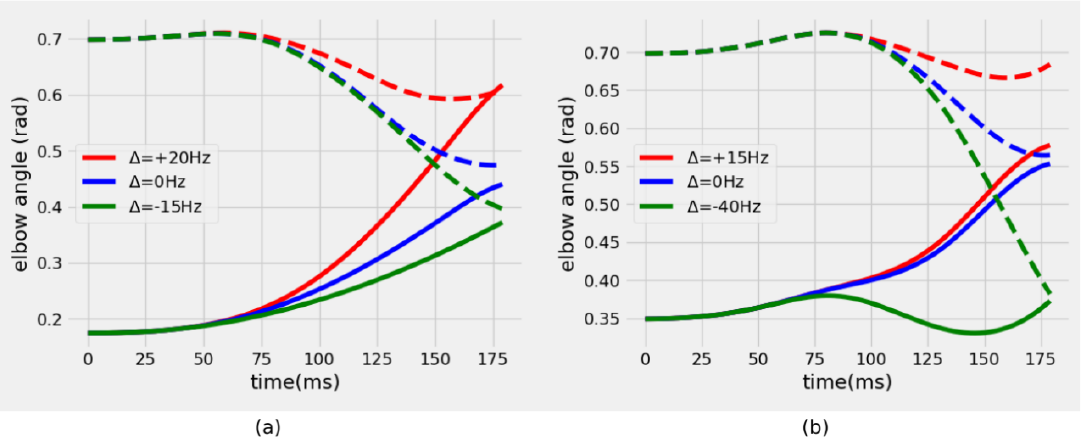
The effect of a constant bias signal (Δ) on the functioning of learnt CMP is shown here. The elbow angle for two different initial angles (solid line (S) and dashed line (D)) for three different bias values can be seen. Changing the amplitude of the constant signal was found to be equivalent to changing the target angle. We observe that the elbow approached the modified target from both the initial angles. (a) is obtained with a simple openloop setup (b) is obtained with the closeloop setup.

### D. Analysis of CMP network

We analyzed the reinforcement learning policy ANN obtained after training. The network received a (1×6) dimensional observation vector and generated (1×2) dimensional action vector (figure 2). There are two fully connected hidden layers each of dimension 64 between the input and output layers. We obtained the activation of each neuron^7^ in the outer hidden layer (*policy net*) and the output layer (*action net*). These activations were obtained for two different initial angles (10 deg and 40 deg) of the elbow joint (figure 11). We observe that the two neurons on the *action net* correlate with the velocity of elbow movement. Presence of such neurons in M1, which encode for velocity have also been observed ([21], [67]).

**Fig. 11.**
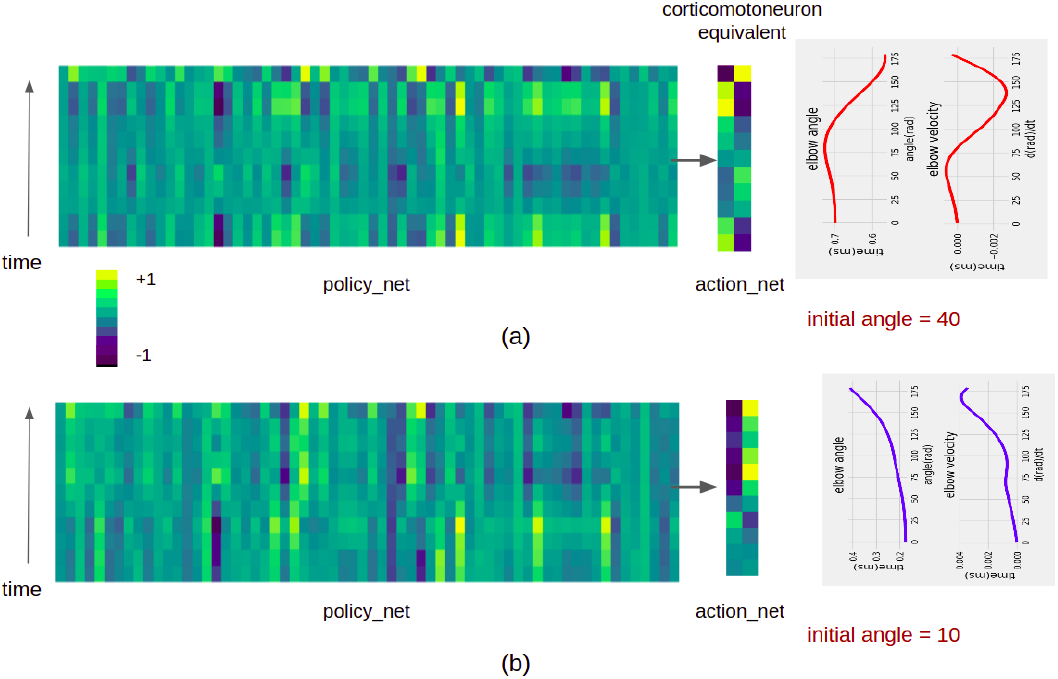
Neuron activation pattern of the 64 neurons in the outer layer of *policy net* and 2 neurons in the outer layer of *action net*. The plot of elbow angle and rate of change of elbow angle is also shown. (a) initial elbow angle = 0.7 rad (b) initial elbow angle = 0.17 rad. The activation pattern of the 2 neurons in the outer layer of *action net* is correlated with rate of change of elbow angle.

The pattern of activation in the neurons of *policy net* was not very obvious. To identify neurons that could potentially be coding for the position or velocity of movement, we defined the metrics (λ_*n*_ and *r_n_*) as follows:

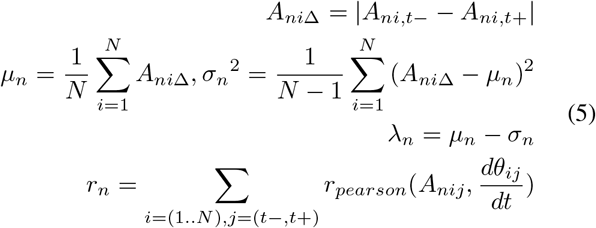

Each experiment involved approaching a specific target angle from two initial angles, one below (*t*−) and one above (*t*+) the target angle (similar to figure 10). *A_ni,t−_* represents the activation of *n^th^* neuron of *policy net* for *i^th^* experiment, with initial angle being below the target angle and *A_ni,t+_* being above the target angle. It was observed that some neurons were found to be within the top-five in all the experiments when the neurons of *policy net* were sorted based on the λ_*n*_ parameter. Figure 12 (a) shows the activation pattern observed for one such neuron (neuron #13). Neurons in the motor cortex have been implicated with such target selections ([68]). The *r_n_* represents the mean pearson correlation coefficient for *n^th^* neuron measured across all experiments. The activation of the neuron is correlated with the rate of change of elbow angle. We find that activation pattern of some neurons (refer figure 12 (b) and (d)) correlate well, as observed in some corticomotor neurons.

**Fig. 12.**
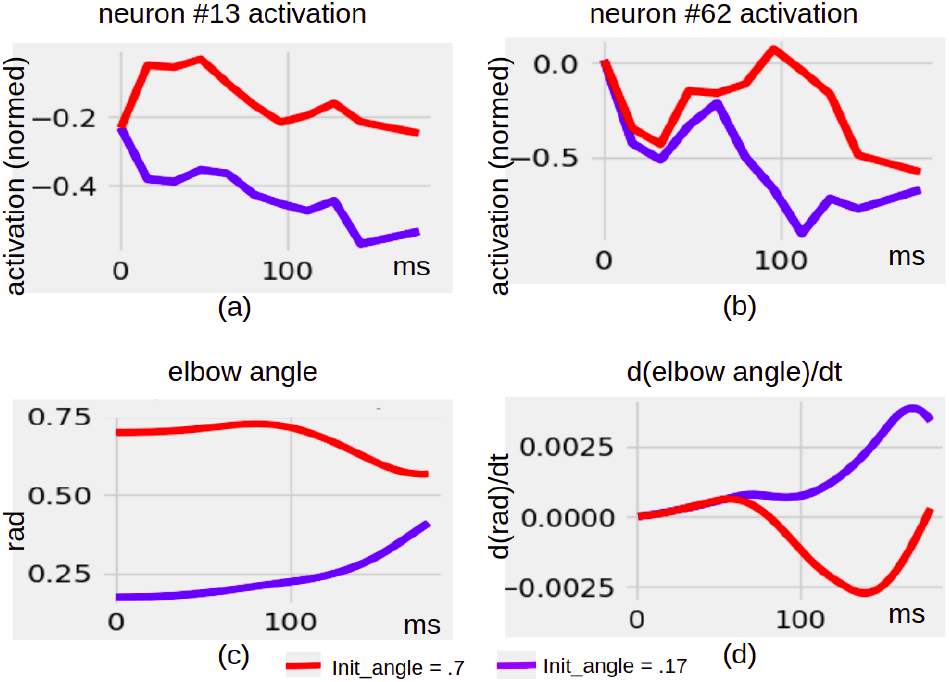
(a) Activation pattern of neuron (#13) in *policy net*. This neuron consistently showed high value for *dm_υ* in all experiments indicating that it could potentially be responsible for trajectory selection. (b) Activation pattern of neuron (#13) in *policy net*. The activation pattern of this neuron showed high correlation with rate of change of elbow angle angle. (c) Plot of elbow angle for two initial angles (0.17 and 0.7 rad) (d) Plot of rate of change of elbow angle for two initial angles (0.17 and 0.7 rad)

Thus, we first built a model that learnt only a single motor program to move the elbow to a fixed target angle from a fixed initial angle. The solution that the RL agent converged to, is one of the many possible solutions for the given problem. Further exploration of the solution space, sensitivity of parameters and other analysis is for a future work. However, it has been shown ([69]–[71]) that such degeneracy is an inherent property of biological systems across scales. We were then able to reach different target positions controlled by a descending drive input. We may extend this and envision a model where the CMP consists of many such submodules (*C*_1_,*C*_2_,…*C_n_*), each representing a different task. Different descending cortical drives (*D_i_*) alter the target for a given task (represented by *C_i_*) to achieve different goals by using the ensemble of 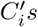. The nervous system is known to demonstrate such compositionality ([66], [72]) to produce complex movements.

## IV. DISCUSSION

In this work, we describe a framework to obtain a functionally equivalent RL policy for the CMP to achieve a specific movement on a virtual physiology consisting of cosimulated neuro-musculoskeletal upperlimb. For the purpose of demonstration in this paper, we chose a single-joint, uni-planar movement. However, the techniques used by us are completely generalizable for more complicated movements involving multiple joints and muscle groups. The platform is modular and hierarchical and allows choosing from a variety of musculoskeletal models and spinal neuronal circuit models independent of each other. The model development involves building a composite model of relevant spinal segments along with their spinal circuitry and the musculoskeletal components. Once the observation and action space are identified for the neuro-musculoskeletal model, the framework described in this paper can be used to learn an artificial neural network equivalent to the CMP. This ANN delivers the patterned stimulations necessary to achieve the specific movement. In the current work, the equivalence and physiological plausibility of the network is achieved by constraining the observation space to the afferent signals from spindle and action space to the activations provided to the spinal networks. We demonstrated that using a two layered network as the RL policy, the learned CMP elicited activations resulting in a muscle activation pattern that mimic the physiologically known triphasic response (section III-B) ([63], [73]). The first agonist burst is thought to initiate the movement (‘‘action burst”), while the antagonist burst is thought to help in halting the movement (“braking pulse”), and the second agonist burst is thought to stabilize the movement (“clamping pulse”) ([73]). Models are known to converge to this result when optimized to have minimum movement time ([74]), minimum torque change ([75]), minimum square magnitude of jerk ([76]). Further, with increasing velocity of movement, the amplitude of the bursts also increased, consistent with recorded observations ([63], [64]). The triphasic muscle response is thought to be due to a central motor program possibly with cortical involvement ([54]).

Descending connections from primary motor cortex are known to terminate on sets of spinal motor neurons. Further cortical microstimulation of primary motor cortex neurons are known to evoke movements to a fixed target area from a variety of initial positions ([66]). These results lead us to hypothesize that our learnt CMP may also be playing the role of parts of the primary motor cortex and their descending cortical signals along with that of spinal interneuronal circuits. The task to move the elbow joint towards a specific invariant position from a range of initial positions can be considered as one of the possible motor program that results in a convergence to an invariant final position. Similarly our behavioural repertoire is known to have numerous such manoeuvres as evidenced in daily life and also by intracortical microstimulation studies ([77], [78]). However it has been suggested that the number of actual synergies implemented are only a handful ([72]).

Our work has demonstrated a possible mechanism by which a rich repertoire of movements can be achieved using a handful of motor programs. In section III-C, we showed that a small injected synaptic current by way of descending cortical fibres modulates the CMP and translates the coordinates of the observation space as seen by the CMP. This results in a shift of the invariant target position. This posits a simple and biologically plausible mechanism by which the same CMP can lead to a family of manoeuvres to different targets. A generalization of this result follows that, by means of a continuously varying current injection, the target may be continuously varied to trace a trajectory consisting of a sequence of targets.

Our work presents an instance of biomimetic movement on a co-simulated neuro-musculoskeletal limb. While biomimetic movement has been achieved earlier in custom neuromuscular models, building virtual movement physiology at scale demands a modular design and the ability to utilize the best advances in the diverse area of neuronal and musculoskeletal modeling. We hope that our work presents a small first step in that direction. We hope to extend our work by realizing the CMP network using models of biological neurons and possibly derive them too using natural learning mechanisms such as STDP, LTP and LTD. Further extensions may be in order to achieve movement to arbitrary target locations in the 3D extrapersonal space. Similarly, adding models for the upstream layers in the form of pre-motor cortices, basal ganglia or cerebellum paves the way for building larger biomimetic systems, one step at a time. Such an in-silico movement physiology platform can play a significant role in realizing the benefits in the form of accelerated development of diagnostic, therapeutic, prosthetic medical devices for motor dysfunctions.

## V. Acknowledgement

The authors would like to gratefully acknowledge the support by way of grant-in-aid provided by the Ministry of Electronics and Information Technology (MEITy-4(11)/2018-ITEA), Government of India.

## Footnotes

1 https://www.aicrowd.com/challenges/neurips-2019-learn-to-move-walkaround

2 http://osim-rl.stanford.edu/docs/nips2017/solutions/

3 http://gym.openai.com/

4 https://github.com/openai/gym/blob/master/docs/creatingenvironments.md

5 https://openai.com/blog/openai-baselines-ppo/

6 https://stable-baselines3.readthedocs.io/

7 https://captum.ai/docs/algorithms#layer-activation

## References

[1] Courtine, G., Gerasimenko, Y., Van Den Brand, R., Yew, A., Musienko, P., Zhong, H.,…Edgerton, V. R. (2009). Transformation of nonfunctional spinal circuits into functional states after the loss of brain input. Nature Neuroscience, 12(10), 1333–1342. https://doi.org/10.1038/NN.2401

[2] Georgopoulos, A. P., & Grillner, S. (1989). Visuomotor coordination in reaching and locomotion. Science (New York, N.Y.), 245(4923), 1209–1210. https://doi.org/10.1126/SCIENCE.2675307

[3] Drew, T., Andujar, J. E., Lajoie, K., & Yakovenko, S. (2008). Cortical mechanisms involved in visuomotor coordination during precision walking. Brain Research Reviews, 57(1), 199–211. https://doi.org/10.1016/J.BRAINRESREV.2007.07.017

[4] Grillner, S. (2018). Evolution: Vertebrate Limb Control over 420 Million Years. Current Biology: CB, 28(4), R162–R164. https://doi.org/10.1016/J.CUB.2017.12.040

[5] Grillner, S., & El Manira, A. (2020). Current Principles of Motor Control, with Special Reference to Vertebrate Locomotion. Physiological Reviews, 100(1), 271–320. https://doi.org/10.1152/PHYSREV.00015.2019

[6] Cheney, P. D., Kasser, R., & Holsapple, J. (1982). Reciprocal effect of single corticomotoneuronal cells on wrist extensor and flexor muscle activity in the primate. Brain Research, 247(1), 164–168. https://doi.org/10.1016/0006-8993(82)91043-5

[7] Kasser, R. J., & Cheney, P. D. (1985). Characteristics of corticomotoneuronal postspike facilitation and reciprocal suppression of EMG activity in the monkey. Journal of Neurophysiology, 53(4), 959–978. https://doi.org/10.1152/JN.1985.53.4.959

[8] Jankowska, E., Padel, Y., & Tanaka, R. (1976). Disynaptic inhibition of spinal motoneurones from the motor cortex in the monkey. The Journal of Physiology, 258(2), 467–487. https://doi.org/10.1113/JPHYSIOL.1976.SP011431

[9] El Manira A, Pombal MA, Grillner S. (1997). Diencephalic projection to reticulospinal neurons involved in the initiation of locomotion in adult lampreys Lampetra fluviatilis. J Comp Neurol. 1997 Dec 29;389(4):603–16. PMID: 9421142.

[10] Pierrot-Deseilligny, E., & Burke, D. (2005). The circuitry of the human spinal cord: Its role in motor control and movement disorders. The Circuitry of the Human Spinal Cord: Its Role in Motor Control and Movement Disorders, 1–642. https://doi.org/10.1017/CBO9780511545047

[11] Kozlov, A., Huss, M., Lansner, A., Kotaleski, J. H., & Grillner, S. (2009). Simple cellular and network control principles govern complex patterns of motor behavior. Proceedings of the National Academy of Sciences of the United States of America, 106(47), 20027–20032.

[12] Kozlov, A. K., Kardamakis, A. A., Kotaleski, J. H., & Grillner, S. (2014). Gating of steering signals through phasic modulation of reticulospinal neurons during locomotion. Proceedings of the National Academy of Sciences of the United States of America, 111(9), 3591–3596.

[13] Talpalar, A. E., Endo, T., Löw, P., Borgius, L., Hägglund, M., Dougherty, K. J.,…Kiehn, O. (2011). Identification of Minimal Neuronal Networks Involved in Flexor-Extensor Alternation in the Mammalian Spinal Cord. Neuron, 71(6), 1071–1084.

[14] Butt, S. J. B., & Kiehn, O. (2003). Functional identification of interneurons responsible for left-right coordination of hindlimbs in mammals. Neuron, 38(6), 953–963.

[15] Kiehn, O. (2016). Decoding the organization of spinal circuits that control locomotion. Nature Reviews Neuroscience 2016 17:4, 17(4), 224–238. https://doi.org/10.1038/nrn.2016.9

[16] Prochazka A. Quantifying proprioception. Prog Brain Res. 1999;123:133–42. PMID: 10635710.

[17] Stienen, A. H. A., Schouten, A. C., Schuurmans, J., & van der Helm, F. C. T. (2007). Analysis of reflex modulation with a biologically realistic neural network. Journal of Computational Neuroscience, 23(3), 333. https://doi.org/10.1007/S10827-007-0037-7

[18] Georgopoulos, A. P., Kalaska, J. F., Caminiti, R., & Massey, J. T. (1982). On the relations between the direction of two-dimensional arm movements and cell discharge in primate motor cortex. The Journal of Neuroscience: The Official Journal of the Society for Neuroscience, 2(11), 1527–1537. https://doi.org/10.1523/JNEUROSCI.02-11-01527.1982

[19] Omrani, M., Kaufman, M. T., Hatsopoulos, N. G., & Cheney, P. D. (2017). Perspectives on classical controversies about the motor cortex. Journal of Neurophysiology, 118(3), 1828–1848. https://doi.org/10.1152/JN.00795.2016

[20] Aflalo, T. N., & Graziano, M. S. A. (2006). Partial tuning of motor cortex neurons to final posture in a free-moving paradigm. Proceedings of the National Academy of Sciences of the United States of America, 103(8), 2909–2914. https://doi.org/10.1073/PNAS.0511139103

[21] Moran, D. W., & Schwartz, A. B. (1999). Motor cortical representation of speed and direction during reaching. Journal of Neurophysiology, 82(5), 2676–2692. https://doi.org/10.1152/JN.1999.82.5.2676

[22] Ashe, J., & Georgopoulos, A. P. (1994). Movement parameters and neural activity in motor cortex and area 5. Cerebral Cortex (New York, N.Y.: 1991), 4(6), 590–600. https://doi.org/10.1093/CERCOR/4.6.590

[23] Smith, A. M., Hepp-Reymond, M. C., & Wyss, U. R. (1975). Relation of activity in precentral cortical neurons to force and rate of force change during isometric contractions of finger muscles. Experimental Brain Research, 23(3), 315–332. https://doi.org/10.1007/BF00239743

[24] Cheney, P. D., & Fetz, E. E. (1980). Functional classes of primate corticomotoneuronal cells and their relation to active force. Journal of Neurophysiology, 44(4), 773–791. https://doi.org/10.1152/JN.1980.44.4.773

[25] Griffin, D. M., Hudson, H. M., Belhaj-Saïf, A., McKiernan, B. J., & Cheney, P. D. (2008). Do corticomotoneuronal cells predict target muscle EMG activity? Journal of Neurophysiology, 99(3), 1169–1186. https://doi.org/10.1152/JN.00906.2007

[26] Hatsopoulos, N. G., Xu, Q., & Amit, Y. (2007). Encoding of Movement Fragments in the Motor Cortex. The Journal of Neuroscience, 27(19), 5105. https://doi.org/10.1523/JNEUROSCI.3570-06.2007

[27] Hatsopoulos, N. G., & Amit, Y. (2012). Synthesizing complex movement fragment representations from motor cortical ensembles. Journal of Physiology, Paris, 106(3–4), 112. https://doi.org/10.1016/J.JPHYSPARIS.2011.09.003

[28] Iyengar, R. S., Pithapuram, M. V., Singh, A. K., & Raghavan, M. (2019). Curated Model Development Using NEUROiD: A Web-Based NEUROmotor Integration and Design Platform. Frontiers in Neuroinformatics, 13, 56. https://doi.org/10.3389/FNINF.2019.00056/BIBTEX

[29] Sutton RS, Barto AG. Introduction to Reinforcement Learning. 1st ed. Cambridge (MA): MIT Press; 1998.

[30] Gottlieb, G. L. (1998). Muscle activation patterns during two types of voluntary single-joint movement. Journal of Neurophysiology, 80(4), 1860–1867. https://doi.org/10.1152/JN.1998.80.4.1860

[31] Morrison, S., & Anson, J. G. (1999). Natural goal-directed movements and the triphasic EMG. Motor Control, 3(4), 346–371. https://doi.org/10.1123/MCJ.3.4.346

[32] Capogrosso, M., Wenger, N., Raspopovic, S., Musienko, P., Beauparlant, J., Luciani, L. B.,…Micera, S. (2013). A Computational Model for Epidural Electrical Stimulation of Spinal Sensorimotor Circuits. Journal of Neuroscience, 33(49), 19326–19340. https://doi.org/10.1523/JNEUROSCI.1688-13.2013

[33] Rybak, I. A., Stecina, K., Shevtsova, N. A., & McCrea, D. A. (2006). Modelling spinal circuitry involved in locomotor pattern generation: insights from the effects of afferent stimulation. The Journal of Physiology, 577(Pt 2), 641. https://doi.org/10.1113/JPHYSIOL.2006.118711

[34] Moraud, E. M., Capogrosso, M., Formento, E., Wenger, N., DiGiovanna, J., Courtine, G., & Micera, S. (2016). Mechanisms Underlying the Neuromodulation of Spinal Circuits for Correcting Gait and Balance Deficits after Spinal Cord Injury. Neuron, 89(4), 814–828. https://doi.org/10.1016/J.NEURON.2016.01.009

[35] Millard, M., Uchida, T., Seth, A., & Delp, S. L. (2013). Flexing computational muscle: modeling and simulation of musculo-tendon dynamics. Journal of Biomechanical Engineering, 135(2). https://doi.org/10.1115/1.4023390

[36] Daniel, M., Hornová, J., Doubrava, K., & Tomanová, M. (2017). Biomechanical analysis of local and global strengthening of gluteus medius. Turkish Journal of Physical Medicine and Rehabilitation, 63(3), 283. https://doi.org/10.5606/TFTRD.2017.916

[37] Cisek, P., Grossberg, S., & Bullock, D. (1998). A corticospinal model of reaching and proprioception under multiple task constraints. Journal of Cognitive Neuroscience, 10(4), 425–444. https://doi.org/10.1162/089892998562852

[38] Sreenivasa, M., Ayusawa, K., & Nakamura, Y. (2016). Modeling and Identification of a Realistic Spiking Neural Network and Musculoskeletal Model of the Human Arm, and an Application to the Stretch Reflex. IEEE Transactions on Neural Systems and Rehabilitation Engineering, 24(5), 591–602. https://doi.org/10.1109/TNSRE.2015.2478858

[39] Tsianos, G. A., Goodner, J., & Loeb, G. E. (2014). Useful properties of spinal circuits for learning and performing planar reaches. Journal of Neural Engineering, 11(5). https://doi.org/10.1088/1741-2560/11/5/056006

[40] Raphael, G., Tsianos, G. A., & Loeb, G. E. (2010). Spinal-Like Regulator Facilitates Control of a Two-Degree-of-Freedom Wrist. Journal of Neuroscience, 30(28), 9431–9444. https://doi.org/10.1523/JNEUROSCI.5537-09.2010

[41] Markin, S. N., Klishko, A. N., Shevtsova, N. A., Lemay, M. A., Prilutsky, B. I., & Rybak, I. A. (2010). Afferent control of locomotor CPG: insights from a simple neuromechanical model. Annals of the New York Academy of Sciences, 1198, 21. https://doi.org/10.1111/J.1749-6632.2010.05435.X

[42] Markin, S. N., Klishko, A. N., Shevtsova, N. A., Lemay, M. A., Prilutsky, B. I., & Rybak, I. A. (2016). A Neuromechanical Model of Spinal Control of Locomotion. 21–65.

[43] Cofer, D., G. Cymbalyuk, J. Reid, Y. Zhu, W. Heitler, & D.H. Edwards, AnimatLab: A 3-D graphics environment for neuromechanical simulations. J Neuroscience Methods., 2010. 187(2): p. 280–288.

[44] Hines, M. L., & Carnevale, N. T. (1997). The NEURON simulation environment. Neural Computation, 9(6), 1179–1209. https://doi.org/10.1162/NECO.1997.9.6.1179

[45] Seth, A., Hicks, J. L., Uchida, T. K., Habib, A., Dembia, C. L., Dunne, J. J.,…Delp, S. L. (2018). OpenSim: Simulating musculoskeletal dynamics and neuromuscular control to study human and animal movement. PLOS Computational Biology, 14(7), e1006223.

[46] Iyengar, R. S., & Raghavan, M. (2020). MPI parallelization of neuroid models using docker swarm. Proceedings of the International Conference on Parallel and Distributed Systems - ICPADS, 2020-December, 655–660. https://doi.org/10.1109/ICPADS51040.2020.00092

[47] Holzbaur, K. R. S., Murray, W. M., & Delp, S. L. (2005). A model of the upper extremity for simulating musculoskeletal surgery and analyzing neuromuscular control. Annals of Biomedical Engineering, 33(6), 829–840. https://doi.org/10.1007/S10439-005-3320-7

[48] Mallampalli, K., Pithapuram, M. V., Rangayyan, Y. M., Iyengar, R. S., Singh, A. K., Sripada, S., & Raghavan, M. (2021). Neuro-musculoskeletal Upper Limb in-silico as virtual patient. BioRxiv, 2021.05.16.444298. https://doi.org/10.1101/2021.05.16.444298

[49] Sengul, Gulgun (2013). Atlas of the Spinal Cord of the Rat, Mouse, Marmoset, Rhesus, and Human. London; Waltham, MA:Academic Press, 2013.

[50] Ko, H. Y., Park, J. H., Shin, Y. B., & Baek, S. Y. (2004). Gross quantitative measurements of spinal cord segments in human. Spinal Cord 2003 42:1, 42(1), 35–40. https://doi.org/10.1038/sj.sc.3101538

[51] Cisi, R. R. L., & Kohn, A. F. (2008). Simulation system of spinal cord motor nuclei and associated nerves and muscles, in a Web-based architecture. Journal of Computational Neuroscience, 25(3), 520–542. https://doi.org/10.1007/S10827-008-0092-8

[52] McIntyre, C. C., & Grill, W. M. (2002). Extracellular stimulation of central neurons: Influence of stimulus waveform and frequency on neuronal output. Journal of Neurophysiology, 88(4), 1592–1604.

[53] Caligiore, D., Pezzulo, G., Baldassarre, G., Bostan, A. C., Strick, P. L., Doya, K., Helmich, R. C., Dirkx, M., Houk, J., Jörntell, H., Lago-Rodriguez, A., Galea, J. M., Miall, R. C., Popa, T., Kishore, A., Verschure, P. F. M. J., Zucca, R., & Herreros, I. (2017). Consensus Paper: Towards a Systems-Level View of Cerebellar Function: the Interplay Between Cerebellum, Basal Ganglia, and Cortex. Cerebellum (London, England), 16(1), 203. https://doi.org/10.1007/S12311-016-0763-3

[54] Graziano, M. S. A., Patel, K. T., & Taylor, C. S. R. (2004). Mapping from motor cortex to biceps and triceps altered by elbow angle. Journal of Neurophysiology, 92(1), 395–407. https://doi.org/10.1152/JN.01241.2003

[55] Todorov, D. I., Capps, R. A., Barnett, W. H., Latash, E. M., Kim, T., Hamade, K. C., Markin, S. N., Rybak, I. A., & Molkov, Y. I. (2019). The interplay between cerebellum and basal ganglia in motor adaptation: A modeling study. PLoS ONE, 14(4). https://doi.org/10.1371/JOURNAL.PONE.0214926

[56] Freeman, S. R., & Durfee, W. K. (2006). Twitch response of intact human tibialis anterior muscle to doublet stimulation at graded strengths. Conference Proceedings: Annual International Conference of the IEEE Engineering in Medicine and Biology Society. IEEE Engineering in Medicine and Biology Society. Annual Conference, Suppl, 6757–6760. https://doi.org/10.1109/IEMBS.2006.260940

[57] Schulman, J., Levine, S., Abbeel, P., Jordan, M.I., & Moritz, P. (2015). Trust Region Policy Optimization. CoRR, abs/1502.05477.

[58] Mnih, V., Badia, A.P., Mirza, M., Graves, A., Lillicrap, T.P., Harley, T., Silver, D., & Kavukcuoglu, K. (2016). Asynchronous Methods for Deep Reinforcement Learning. Proceedings of The 33rd International Conference on Machine Learning volume 48 of Proceedings of Machine Learning Research, pp. 1928–1937, 20–22 Jun 2016‥

[59] Wang, Z., Bapst, V., Heess, N., Mnih, V., Munos, R., Kavukcuoglu, K., de Freitas, N. (2017): Sample Efficient Actor-Critic with Experience Replay. In: ICLR17

[60] Schulman, J., Moritz, P., Levine, S., Jordan, M.I., & Abbeel, P. (2016). High-Dimensional Continuous Control Using Generalized Advantage Estimation. CoRR, abs/1506.02438.

[61] Bengio, Y., Louradour, J., Collobert, R., & Weston, J. (2009). Curriculum learning. ACM International Conference Proceeding Series, 382. https://doi.org/10.1145/1553374.1553380

[62] Fuglevand, A. J., Winter, D. A., & Patla, A. E. (1993). Models of recruitment and rate coding organization in motorunit pools. Journal of Neurophysiology, 70(6), 2470–2488. https://doi.org/10.1152/JN.1993.70.6.2470

[63] Hallett, M., & Marsden, C. D. (1979). Ballistic flexion movements of the human thumb. The Journal of Physiology, 294(1), 33. https://doi.org/10.1113/JPHYSIOL.1979.SP012913

[64] Mustard, B. E., & Lee, R. G. (1987). Relationship between EMG patterns and kinematic properties for flexion movements at the human wrist. Experimental Brain Research, 66(2), 247–256. https://doi.org/10.1007/BF00243302

[65] Loeb, G. E. (1983). Finding common groud between robotics and physiology. In Trends in Neurosciences (Vol. 6, Issue C, pp. 203–204). https://doi.org/10.1016/0166-2236(83)90093-0

[66] Graziano, M. S. A., Taylor, C. S. R., & Moore, T. (2002). Complex movements evoked by microstimulation of precentral cortex. Neuron, 34(5), 841–851. https://doi.org/10.1016/S0896-6273(02)00698-0

[67] . Paninski L, Fellows MR, Hatsopoulos NG, Donoghue JP (2004). Spatiotemporal tuning of motor cortical neurons for hand position and velocity. J Neurophysiol 91: 515–532, 2004.

[68] Omrani M, Murnaghan CD, Pruszynski JA, Scott SH (2016). Distributed task-specific processing of somatosensory feedback for voluntary motor control. eLife 5: e13141, 2016.

[69] Prinz, A. A., Bucher, D., & Marder, E. (2004). Similar network activity from disparate circuit parameters. Nature Neuroscience 2004 7:12, 7(12), 1345–1352. https://doi.org/10.1038/nn1352

[70] Edelman, G. M., & Gally, J. A. (2001). Degeneracy and complexity in biological systems. Proceedings of the National Academy of Sciences of the United States of America, 98(24), 13763–13768. https://doi.org/10.1073/PNAS.231499798

[71] Rathour, R. K., & Narayanan, R. (2014). Homeostasis of functional maps in active dendrites emerges in the absence of individual channelostasis. Proceedings of the National Academy of Sciences of the United States of America, 111(17).

[72] Bizzi, E., Mussa-Ivaldi, F. A., & Giszter, S. (1991). Computations underlying the execution of movement: a biological perspective. Science (New York, N.Y.), 253(5017), 287–291. https://doi.org/10.1126/SCIENCE.1857964

[73] Hannaford, B., Cheron, G., & Stark, L. (1985). Effects of applied vibration on triphasic electromyographic patterns in neurologically ballistic head movements. Experimental Neurology, 88(2), 447–460. https://doi.org/10.1016/0014-4886(85)90206-7

[74] Hatze, H. (1976). The complete optimization of a human motion. Mathematical Biosciences, 28(1–2), 99–135. https://doi.org/10.1016/0025-5564(76)90098-5

[75] Uno, Y., Kawato, M., & Suzuki, R. (1989). Formation and control of optimal trajectory in human multijoint arm movement. Biological Cybernetics 1989 61:2, 61(2), 89–101. https://doi.org/10.1007/BF00204593

[76] Flash, T., & Hogan, N. (1985). The coordination of arm movements: an experimentally confirmed mathematical model. The Journal of Neuroscience: The Official Journal of the Society for Neuroscience, 5(7), 1688–1703. https://doi.org/10.1523/JNEUROSCI.05-07-01688.1985

[77] Graziano, M. (2006). The organization of behavioral repertoire in motor cortex. Annual Review of Neuroscience, 29, 105–134.

[78] Graziano, M. S. A. (2016). Ethological Action Maps: A Paradigm Shift for the Motor Cortex. Trends in Cognitive Sciences, 20(2), 121–132.

